# KDM5 histone-demethylases contribute to replication stress response and tolerance

**DOI:** 10.1101/2019.12.16.877399

**Authors:** Solenne Gaillard, Virginie Charasson, Cyril Ribeyre, Kader Salifou, Marie-Jeanne Pillaire, Jean-Sebastien Hoffmann, Angelos Constantinou, Didier Trouche, Marie Vandromme

## Abstract

KDM5A and KDM5B histone-demethylases are overexpressed in many cancers and have been involved in drug tolerance. Here, we describe that KDM5A, together with KDM5B, contribute to replication stress (RS) response and tolerance. First, they positively regulate RRM2, the regulatory subunit of Ribonucleotide Reductase. Second, they are required for optimal activation of Chk1, a major player of the intra-S phase checkpoint that protects cells from RS. This role in Chk1 activation is probably direct since KDM5A is enriched at ongoing replication forks and associates with both PCNA and Chk1. Because RRM2 is a major determinant of replication stress tolerance, we developed cells resistant to HU, and show that KDM5A/B proteins are required for both RRM2 overexpression and tolerance to HU, in a manner that is independent of their demethylase activity. Altogether, our results indicate that KDM5A/B are major players of RS management. They also show that drugs targeting the enzymatic activity of KDM5 proteins may not affect all cancer-related consequences of KDM5A/B overexpression.

## INTRODUCTION

KDM5 proteins belong to the JUMONJI family of histone demethylases that catalyze the demethylation of di- and tri-methylated lysines on histone and non-histone proteins. KDM5A, also known as JARID1A or RBP2 (Retinoblastoma binding protein 2), was originally discovered as a Retinoblastoma binding protein [1]. Further studies showed that KDM5A is a histone-demethylase specific to the di- and tri-methylated forms of Lys4 of Histone H3 (H3K4me2/3), a mark associated with promoters of transcriptionally active genes [2–4]. In agreement with its demethylase activity against an active mark of transcription, KDM5A is involved in gene repression by demethylating H3K4me3 at gene promoters. It participates in many repressive chromatin complexes, including the Polycomb complex [5], the Sin3 corepressor complex [6] and the NuRD complex [7]. KDM5A was involved in the stable repression of E2F dependent genes that occurs during terminal differentiation and senescence, two biological processes requiring permanent exit from the cell cycle [6, 8]. Rb that is up-regulated during differentiation was shown to sequestrate KDM5A/Rbp2, thus impeding its repressor activity on genes required for differentiation [9]. Whereas its role in the stable repression of E2F dependent genes has been well documented, whether it is also important for the regulation of the oscillation of E2F genes transcription during the cell cycle, is less known. However, a recent report describes its interaction with p130 in order to demethylate H3K4 on E2F promoters in G0 and early G1 [10].

KDM5A can also act as a transcriptional activator. When enriched in gene bodies, it plays a role in the elongation step of Polymerase II (Pol-II) by maintaining low levels of H3K4 methylation [11]. In addition, KDM5A activates genes critical for mitochondrial function and metabolism, in a manner that is independent of its demethylase activity but dependent of its C-terminal PHD3 domain that serves as docking site to H3K4me3 at promoters [12]. Interestingly, genome-wide approaches revealed that KDM5A is enriched at promoters of highly expressed genes marked by H3K4me3 [12–14], suggesting that its demethylase activity is tightly regulated, or targets substrates other than histones. Thus, KDM5A plays both positive and negative roles on transcription, in a manner that is either dependent or independent of its demethylase activity against H3K4me2/3, and serves as a platform to recruit other chromatin regulators [3, 7, 15]. This latter property is well illustrated by its role in homologous recombination repair of DNA double strand breaks (DSB), in which it is needed to recruit to DSB the ZMYND8-NurD complex [16].

Noticeably, KDM5A was recently identified as a key factor for drug tolerance in different models of cancer. This property was first described in the non-small cell lung cancer PC9, carrying a mutation in EGFR receptor. When exposed to Erlotinib, an EGF receptor inhibitor, a small percentage of cells, called Drug Tolerant Persister Cells (DTPs) become tolerant to this drug, in a manner that depends on KDM5A overexpression and activity. These cells then start to proliferate in the presence of erlotinib and are called Drug Tolerant Expanded Persisters Cells (DTEPs) [17]. It was further shown that a similar mechanism is involved in the emergence of tolerance to drug treatment in distinct types of cancer cells. This is true both for drugs targetting specific molecules or functioning as cytotoxic agents such as Cisplatin[18]. A specific inhibitor of KDM5A demethylase activity, CPI-455, was shown to impede the emergence of the so-called DPC cells in various cancer cell lines treated with different chemotherapeutic drugs. CPI-455 inhibits to similar extents KDM5A, B and C [18]. In agreement with a key role for KDM5 histone-demethylases in cancer progression and drug tolerance, KDM5A and KDM5B are up-regulated in many cancers, including gastric, breast and lung cancers as well as acute myeloid leukemia reviewed in [19, 20]. KDM5A or KDM5B up-regulation appears as a key event in the development of drug tolerance in cancer cells [18, 21] [22, 23].

Replication stress (RS), referring to any hindrance to progression of the replication fork during S-phase, has been proposed to be an early step of carcinogenesis by triggering genome instability. The generation of aberrant replication fork structures containing single stranded DNA activates the RS response, primarily mediated by the kinase ATR (ATM-­- and Rad3-­-related) and its downstream effector, Chk1. This results in S-­-phase arrest in order to allow replication stress resolution and fork restart. This mechanism ensures that the DNA is faithfully duplicated, and only once, at each cell cycle [24, 25]. Because of their high proliferation rate, cancer cells harbour a high RS and develop strategies to tolerate this level of RS, such as overexpression of RRM2, a subunit of the Ribonucleotide Reductase (RNR), that catalyzes the formation of deoxyribonucleotides from ribonucleotides, and thus is involved in the supply of dNTPs during S-phase [26].

Here we describe a role for KDM5A in managing the replication stress response by fine-­-tuning the expression of RRM2 in normal and stressed conditions, and by regulating the activation of Chk1 by ATR in response to HU. Importantly, we show that the resistance of cells to permanent exposition to HU depends on KDM5A/B dependent up-­-regulation of RRM2, in a manner that is independent of the demethylase activity.

## RESULTS

### KDM5A and KDM5B positively regulate RRM2

Because KDM5A is involved in the stable repression of E2F-dependent genes during differentiation and senescence, we asked if KDM5A regulates these genes in proliferative U2OS cells. We also analysed the role of KDM5B, since KDM5B can compensate for the absence of KDM5A on a subset of gene promoters [8, 27]. KDM5B or/and KDM5A were depleted by RNA interference using specific siRNA (siK5A, siK5B), and mRNA expression levels of some E2F genes were quantified 48 hours later. A pool of non-targeting siRNA was used as control (siCtle). CDC6, a *bona fide* E2F gene, was not affected. Surprisingly however, two other E2F target genes, CCNE1 and RRM2, decreased about two fold only when KDM5A and B were depleted together (Fig1A). Decreased expression of RRM2 was also observed at the protein level, using two distinct set of siRNA targeting KDM5A and KDM5B (Fig1B). However, despite this effect on gene regulation, and particularly the drop in RRM2 at both the mRNA and protein levels, the cell cycle of U2OS cells was not disturbed (Fig 1C) and cell survival was only slightly decreased (Fig 1D).

**Figure 1.**
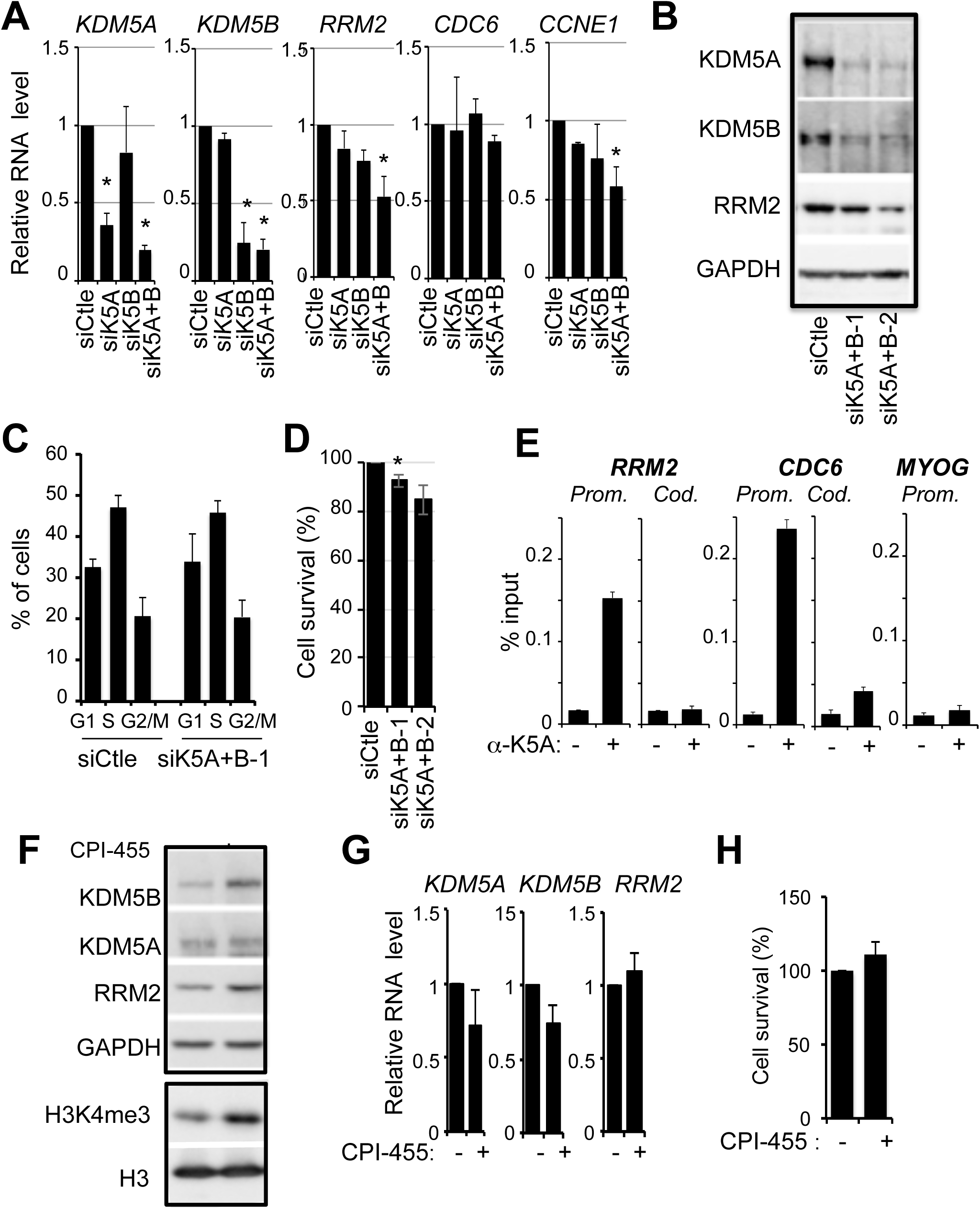
KDM5A/B positively regulates RRM2 expression. **A-** Relative mRNA expression levels of KDM5A, KDM5B, and the E2F target genes RRM2, CDC6 and CCNE1 upon transfection of the indicated siRNA in U2OS cells (siCtle corresponds to a non-targeting pool of siRNA). Expression levels were normalized to the reference gene P0 (ribosomal phosphoprotein P0) and calculated relative to 1 for the siCtle sample. The mean and standard deviation from 3 independent experiments are shown. The star (*) indicates significant difference between the siCtle and the K5A/B siRNA treated cells (pvalue <0.05 calculated using a paired t-test). **B-** Western-blot analysis of KDM5A, KDM5B, RRM2 and GAPDH as a loading control from U2OS cells transfected with siRNA directed against KDM5A and KDM5B. Two distinct couples of siRNA (siK5A+B-1 and -2) were used. **C-** Cell cycle distribution of U2OS cells depleted for KDM5A and KDM5B using siK5A+B-1 compared to siCtle treated cells, analyzed by the high content imaging system Operetta following EdU labeling and DAPI staining. **D-** Percentage of living cells following depletion of KDM5A and B using siK5A+B-1 siRNAs. The mean and standard deviation from 3 independent experiments are shown, following normalization to 100 for siCtle treated cells. Paired t-tests indicate a pvalue <0.05 between the first couple of siK5A/B siRNA and siCtle treated cells (*) but not for the second one with pvalue=0.054. **E-** ChIP analysis of KDM5A on the RRM2 and CDC6 promoter (Prom.) and coding (Cod.) regions. The myogenin gene is not expressed in U2OS cells and its promoter serves as a negative control. A representative experiment out of 4 is shown. **F-** Western-blot analysis of KDM5A, KDM5B, RRM2 and GAPDH from U2OS cells treated each 24 hours or not with KDM5 inhibitor CPI-455 for 48 hours. **G-** Relative mRNA expression levels of KDM5A, KDM5B and RRM2 in cells treated, each 24 hours, with 12.5 mM KDM5 inhibitor CPI-455 (+) or DMSO (-) for 48 hours. Expression levels were normalized to the reference gene P0 (ribosomal phosphoprotein P0) and calculated relative to 1 for the siCtle sample. The mean and standard deviation from 3 independent experiments are shown. A paired t-test indicated no significant difference for all tested genes between CPI treated and untreated cells. **H-** Percentage of living cells following treatment each 24 hours with CPI-455 for 72 hours (+) or DMSO (-). The mean and standard deviation from 3 independent experiments are shown, following normalization to 100 for DMSO treated cells. A paired t-test indicated no significant difference between CPI treated and untreated cells.

We next performed ChIP experiments to investigate whether KDM5A is recruited to the promoter of RRM2, which is regulated by KDM5 proteins, and that of CDC6, which is not. As shown in Fig1E, KDM5A was enriched at the promoter regions of both RRM2 and CDC6, and not at their coding sequences nor at the promoter of a gene inactive in non-muscle cells (MYOG). These results are consistent with genome-wide data showing that KDM5A binds to the promoter of actively transcribed genes but that its loss affects only a subset of genes [28, 29].

Next, we questioned if the demethylase activity of KDM5A/B is required for regulating RRM2 expression. We made use of CPI-455, a well-described and highly specific inhibitor of KDM5 enzymatic activity [18]. U2OS cells were treated for 2 days once a day with 12.5 μM CPI-455. As expected, this treatment led to an increase in H3K4me3 levels (Fig 1F). A slight increase in both KDM5B and RRM2 proteins expression was observed by western-blot, whereas KDM5A expression was unchanged (Fig 1F). However, KDM5A, KDM5B and RRM2 expressions were not changed or weakly affected at the mRNA level (Fig 1G). Thus, RRM2 expression is dependent on KDM5A/B proteins, but not on their demethylase activity. Accordingly, no change in cell survival could be observed upon CPI-455 treatment (Fig 1H).

All together, these data show that KDM5A/B act as a positive regulator of RRM2 probably through direct control of its promoter activity and in a demethylase-independent manner.

### KDM5A/B restrains replication stress in response to HU

RRM2 is a subunit of the Ribonucleotide Reductase (RNR), catalyzing the formation of deoxyribonucleotides from ribonucleotides, and is thus involved in the supply of dNTPs during S-phase. Under RNR inhibition, replication forks stall, leading to replicative stress and ultimately collapse to generate double strand breaks (reviewed in [26, 30]. In regard of its effect on RRM2 expression, we looked if depletion of KDM5A/B could trigger endogenous replication stress. As a read-out, we monitored the consequences this stress could have on fragile sites breakage during S-phase. Indeed, fragile sites are regions of the genome hard to replicate, and thus hypersensitive to replication stress. When incompletely replicated, they are found into 53BP1 bodies in the next G1 phase, waiting for the following S phase to be fully replicated [31–33]. U2OS cells were treated with siRNA against KDM5A/B or a non-targeting siRNA pool as control (siCtle), and allowed to grow for 48 hours. Following a 30 min incubation with EdU, that incorporates into replicating DNA during S-phase, cells were stained for 53BP1, EdU and DNA content using DAPI. Immunofluorescence signals were analyzed using a high content imaging system and quantified using the Columbus software. As illustrated in Fig S1, plotting EdU labeling versus DAPI stain allowed separating cells that were in G1, S or G2/M and quantifying the number of 53BP1 bodies per nucleus in G1 phase. As a control, we treated U2OS cells with a low dose of aphidicolin, a condition known to increase 53BP1 bodies in G1 cells. As expected, the number of G1 53BP1 foci increases upon treatment with aphidicolin (Fig S1). As shown in Figure 2A, the % of G1 cells with a high number of 53BP1 bodies increased in cells depleted for KDM5A/B compared to siCtle treated cells, whereas the number of cells with no or only one 53BP1 body decreased accordingly. Thus, depletion of KDM5A and B leads to replication-stress induced DNA damage, that may be due in part to RRM2 down-regulation.

**Figure 2.**
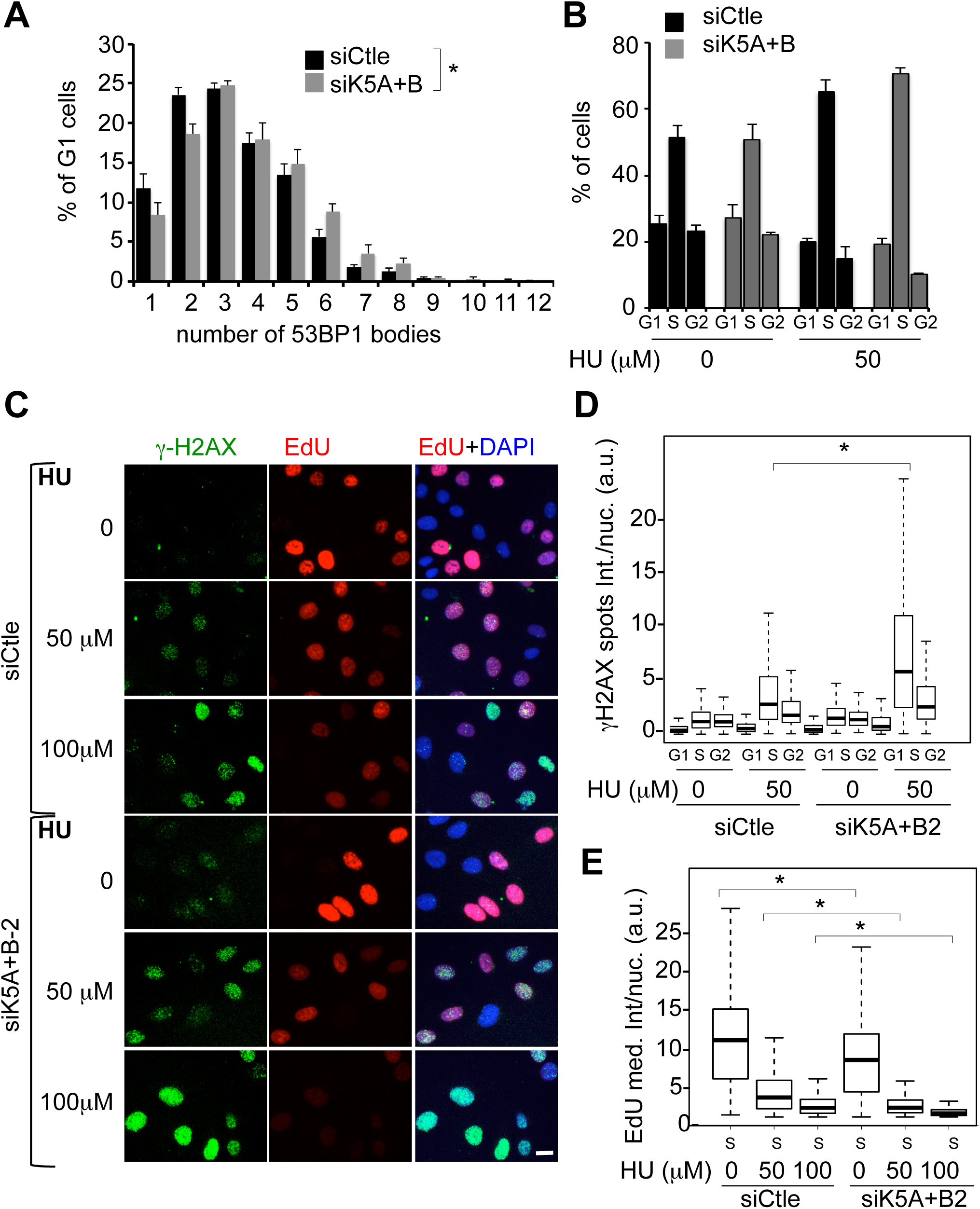
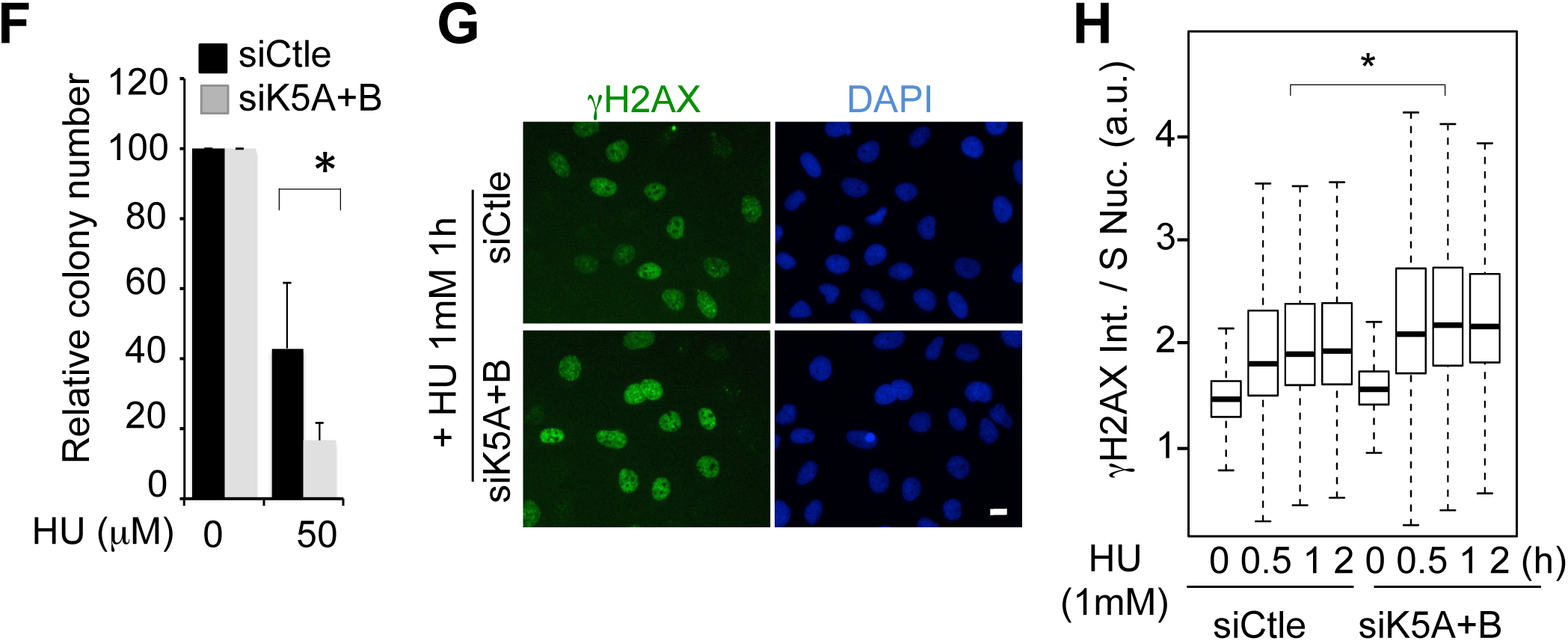
KDM5A/B protect cells from replication stress. **A-** U2OS cells were transfected with with siRNA directed against KDM5A and KDM5B (siK5A+B-1), or a non-targeting siRNA pool as control (siCtle) and stained for 53BP1, EdU (added in the medium 30 min before fixation) and DAPI. Images were acquired with the Operetta device and 53BP1 bodies were counted in G1 cells nuclei (sorted by EdU/DAPI staining), using the Colombus software. See SupFig2 for method illustration. Number of G1 cells analyzed was > 700 for each condition. Difference between siCtle and siK5A/B treated cells was found significant by Wilcoxon (pvalue=1.83*10^-7^). **B-** Cell cycle distribution of U2OS cells treated for 48 hours with the indicated siRNA. U2OS cells were transfected as in A and the last 24 hours incubated with 50 μM HU or left untreated. Cells were labeled with EdU during 30 min before fixation. Cells were stained for EdU and DAPI, and images were acquired using the operetta device. Cell cycle distribution was analyzed thanks to the colombus software. **C-** Cells were transfected as in B, and treated with the indicated doses of HU or left untreated. Cells were labeled with EdU during 30 min before and stained for γ-H2AX, EdU incorporation, and DNA content by DAPI. Images were acquired using the high-content imaging device Operetta. Images of the experiment quantified in Fig2D are shown. Bar=5μM **D-** γH2AX staining was quantified in the nuclei (spots total intensity) of cells, treated or not with 50 μM HU, in G1, S and G2 phases of the cell cycle sorted by EdU/DAPI staining. At 100 μM HU, EdU staining was too low in siKDM5A/B depleted cells to accurately separate G1, S and G2 cells (see panel E). A representative experiment out of 3 is shown. Results are presented as box-­-plot showing the median, the 25 % and 75% quantiles and extrema. Number of counted cells is > 1500 cells for each point. * indicates a pvalue < 1*10^-50^. **E-** Experiment of panels C-D was quantified for EdU in S-phase nuclei. The median intensity per nucleus is represented as a box-plot. Number of S-phase cells > 900 for each point. The star * indicates a significant difference between siCtle and siKDM5A+B-2 treated cells for each dose of HU calculated by Wilcoxon (pvalue=7.92*10^-27^, pvalue= 3.52*10^-66,^ pvalue=1.66*10^-101^ at 0, 50 and 100μM, respectively). **F-** Clonogenic assay of cells treated with siRNA directed against KDM5A-1 and KDM5B-1 or a non-targeting siRNA pool as control (siCtle), and exposed for 24 hours to 50 μM HU, before allowing clones to grow for 10 days more. The mean and standard deviation from 3 independent experiments are shown, following normalization to 100 for untreated cells. The star * indicates a significant difference between siK5A+B and siCtle treated cells with a pvalue < 0.05 (paired t test). **G-** As in B, except that cells were treated for only 0, 0.5, 1 and 2 h with 1mM HU. Cells were stained for γ-H2AX and DNA content by DAPI. Images were acquired using the operetta device. Images of cells treated 1 h with 1mM HU of the experiment quantified in Fig. 2H are shown. Bar=5μM **H-** Total nuclear γ-H2AX was quantified in S-phase nuclei of U2OS cells, transfected with the indicated siRNA, and treated with 1mM HU for the indicated time. S-phase cells were sorted out according to the DAPI staining and results are presented as box-­-plot. A representative experiment out of 3 is shown. Number of S-phase cells examined > 1200 for each point. *A Wilcoxon-test between siK5A/B and siCtle treated cells indicates significant difference at each time point of HU treatment (pvalue < 4.64*10^-45^ at 0.5 h, pvalue = 3.86*10^-32^ at 1 h, pvalue= 1.09*10^-29^at 2 h).

We then investigated whether depletion of KDM5A/B could exacerbate the consequence of induced replication stress. We used Hydroxyurea (HU), a potent inhibitor of RNR, which inhibits DNA replication by triggering an unbalance in the dNTPs pool. We analysed the phosphorylation of H2AX (γ-H2AX), a hallmark of the DNA damage response (DDR). U2OS cells were treated with siRNA against KDM5A/B, allowed to grow for 24 hours, before exposure to 50 μM HU for further 24 hours. Following a 30 min incubation with EdU, that incorporates into replicating DNA during S-phase, cells were stained for γ-H2AX, EdU and DNA content with DAPI. Signals were analyzed using a high content imaging system, and γH2AX spots intensity were quantified in cells in G1, S and G2 phases of the cell cycle. As expected, treatment with HU led to a block in S-phase (Fig 2B), and an increase in nuclear γ-H2AX content in S-phase cells nuclei (Fig 2C-D). Depletion of KDM5A/B did not change the cell cycle distribution upon HU treatment (Fig 2B). Strikingly however, depletion of KDM5A/B further increased the level of γ-H2AX induced by HU in S-phase nuclei compared to control cells (Fig 2C-D), accompanied by a decrease in EdU incorporation in S-phase cells (Fig 2E). Importantly, these results were reproduced with a distinct set of siRNA targeting KDM5A/B, excluding off-target effects (Fig S2). This increase in γ-H2AX coupled to a decrease in DNA synthesis suggests that KDM5A/B protect cells from HU-induced replication stress. To test this possibility, we performed a clonogenic assay. This showed that co-depletion of KDM5A and KDM5B sensitizes cells to HU (Fig 2F), pointing to a role of KDM5A/B in managing the response to replication stress. However, because long term treatment with HU are known to induce fork collapse, and subsequent generation of replication-dependent DNA double strand breaks [34, 35], we could not exclude that the results obtained above reflect the role of KDM5A/B in DNA breaks repair that was recently described [16]. To investigate whether KDM5A/B plays a direct role in the replication stress response besides its function in DSB repair, we looked for γ-H2AX signal intensity in nuclei of cells treated for shorter time with 1 mM HU, conditions known to induce rapid and robust replication stress. At these concentrations of HU, replication is almost stopped and EdU cannot be used to follow the cell cycle distribution. Thus, S-phase cells were sorted out by DAPI stain intensity. As shown in Fig2 G-H, cells depleted for KDM5A/B displayed higher levels of γ-H2AX than control cells.

All together, these results show that endogenous or induced replication stress is enhanced upon depletion of KDM5A/B, suggesting a role of KDM5A/B in protecting the genome from this stress.

### KDM5A/B are required for full activation of Chk1 in response to RS

The response to RS mostly relies on the activation of the sensor kinase ATR, which phosphorylates itself on Thr1989 [36, 37] and the effector kinase Chk1 on Ser345 [38], resulting in its activation. This pathway is quickly activated upon RS, protecting the genome and allowing cells to recover from the stress. The activation of Chk1 by ATR at stalled forks depends on many signalling adapter components such as TopBP1, the rad9-rad1-Hus1 (9-1-1) complex, and Claspin [24, 25, 39]. In order to investigate the consequence of KDM5A/B depletion on replication stress response, we first analysed the expression of the mRNA encoding these proteins by RT-qPCR (Fig 3A and Sup Fig 3). ATR, CHK1, HUS1 and CLASPIN mRNA were decreased upon depletion of KDM5A/B, and the other tested genes were unchanged (Figure 3A and S3), which could result in a defect in the ATR-Chk1 pathway. Note however that no reproducible effect could be seen on ATR and Chk1 protein expression in the unchallenged conditions (Fig 3B-C), suggesting that the decrease of ATR and Chk1 mRNA levels we detect is probably not functionally important.

**Figure 3:**
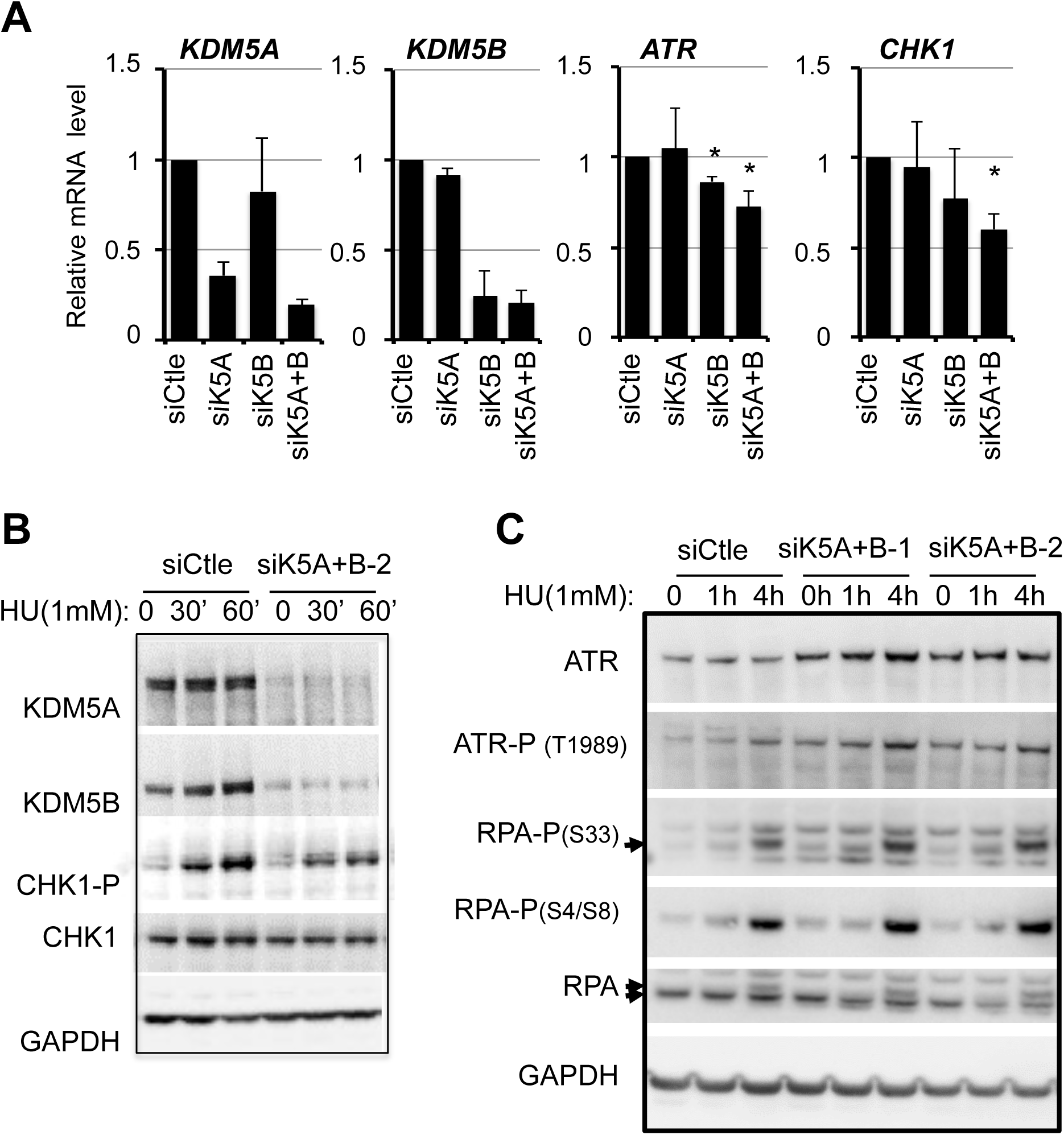
KDM5A/B are involved in Chk1 activation. **A-­-** Relative mRNA expression of KDM5A, KDM5B, ATR and Chk1 in U2OS cells treated with siRNA directed against KDM5A or/and KDM5B or a non-­-targeting siRNA as control (siCtle). mRNA expression is normalized with the reference gene P0, and calculated relative to 1 for the siCtle. The mean and standard deviation from 3 independent experiments are shown. The star (*) indicates a pvalue < 0.05 calculated using a paired t-­-test. **B-­-** U2OS cells were transfected by the indicated siRNA and subjected to western-­-blot analysis of KDM5A, KDM5B, Chk1 and S345 phospho Chk1 (CHK1-­-P). GAPDH is used as a loading control. **C-­-** U2OS cells were transfected by the indicated siRNA and subjected to western-­-blot analysis of KDM5A, KDM5B, Ser1989 phospho ATR (P-­-ATR), S4/S8 phospho RPA (P-­-RPA), ATR and RPA. GAPDH serves as a loading control.

We next analysed the effect of KDM5A depletion on Chk1 pathway activation. Upon treatment with 1mM HU, Chk1 was rapidly activated as revealed by its phosphorylation on Ser345 (Fig 3B). This phosphorylation was seen as soon as 30 min following treatment and is further enhanced after one hour. Upon depletion of KDM5A/B, Ser345 phosphorylation of Chk1 and thus its activation was strongly affected (Fig 3B). Importantly, this was observed using another independent couple of siRNAs directed against KDM5A/B (Fig S4). By contrast, knocking-down KDM5A/B did not impede phosphorylation of ATR on Thr1989 reflecting ATR activation, nor phosphorylation of RPA on two distinct sites, one targeted by ATR (S33) and the other by DNA-PK (S4/S8) (Fig 3C). On the contrary, phosphorylation of ATR and RPA was slightly induced upon KDM5A/B depletion, as shown above for γH2A.X. Thus, these data demonstrate that Chk1 activation is defective in KDM5A/B-deficient cells despite efficient phosphorylation of ATR-Thr1989, phosphorylation of H2A.X (Fig 2C), and RPA-Ser33 (Fig 3C), two other substrates of ATR. Altogether, our results thus indicate that KDM5A/B is specifically required for optimal phosphorylation of Chk1 by ATR.

### KDM5A localizes to the fork and interacts with PCNA

We next investigated the mechanism by which KDM5A/B could participate in Chk1 activation. Activation of Chk1 is mediated by its ATR-dependent phosphorylation at stalled replication forks. Interestingly, KDM5C was shown to localize at replication forks and interact with PCNA [40, 41]. We thus envisioned that KDM5A/B could also localize at replication forks in order to regulate Chk1 activation. To test this possibility, we performed iPOND experiments, which allow to analyse the proteins present at on-going replication forks by western blot. Cells were labelled 5 min with EdU, which incorporates at active replication forks and allows their purification. As controls, EdU was either not coupled with biotin (Click -) or EdU-labelled cells were submitted to a thymidine chase in order to compare replication fork versus fork-free chromatin (Fig4A). As shown in Figure 4B, KDM5A and KDM5B were specifically enriched at on-going replication forks when compared to the thymidine chase condition, as was PCNA used as a positive control. Note however that a significant amount of KDM5A and KDM5B persisted following thymidine chase, indicating that they also have a global chromatin distribution in agreement with their role in transcription regulation. Next, we investigated whether KDM5A/B recruitment at forks is modulated by replication stress, by treating cells with 1mM HU following the 5 min EdU labelling. In those conditions, replication is stopped and iPOND allows the isolation of stalled replication forks (30 to 120 min of 1mM HU, Fig4A). As previously described [42], Rad51 was recruited to stalled forks, whereas PCNA was released (Fig 4B). KDM5A and KDM5B behaved as PCNA and were released from the fork upon replication stress, decreasing to the levels observed upon thymidine chase (Fig. 4B).

**Figure 4:**
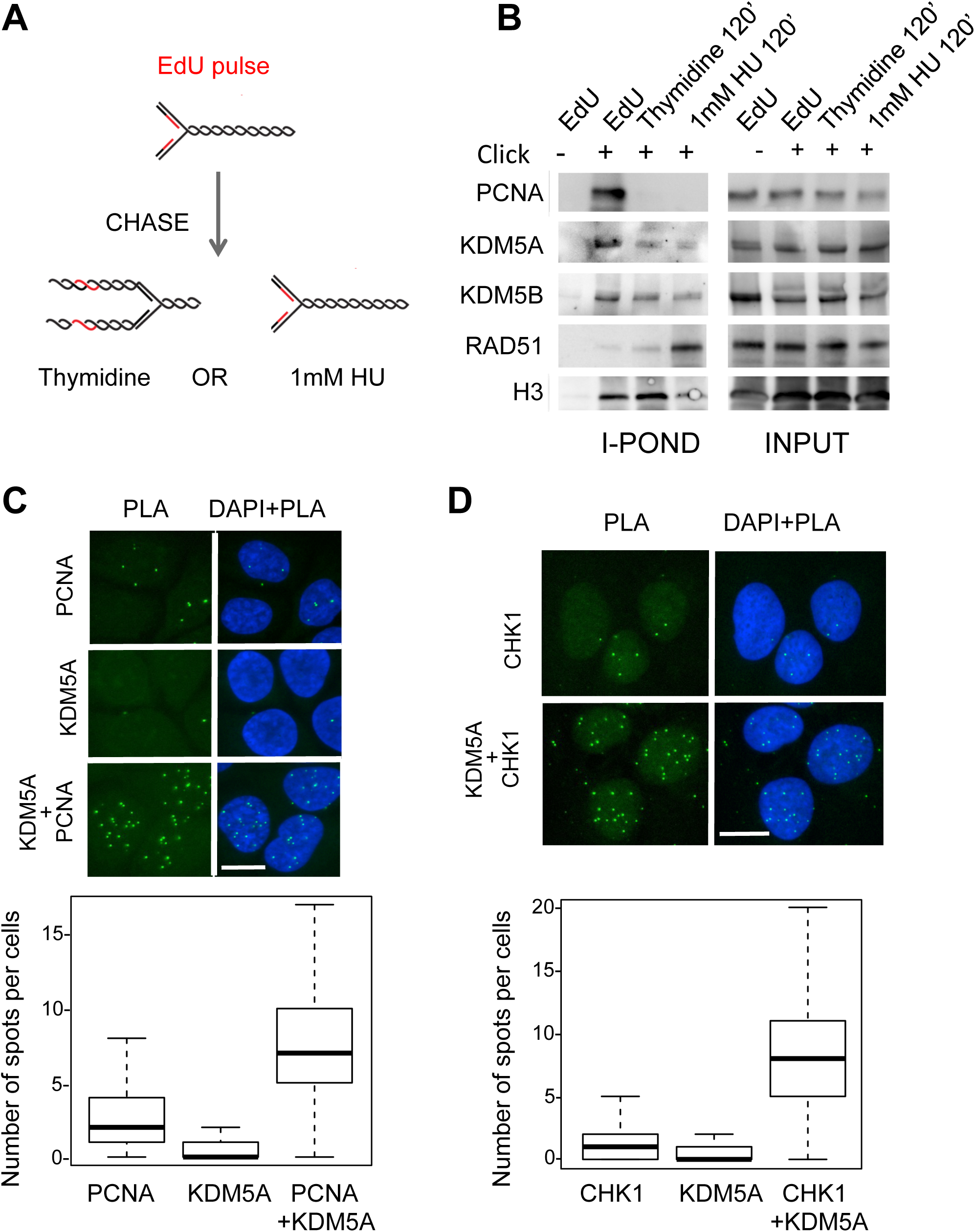
KDM5A is recruited at replication forks in proximity with PCNA and Chk1. **A-­-** Schematic description of the iPOND experiment. Active forks are labeled with EdU. Upon thymidine chase, labeled DNA is not associated with a fork. Upon HU treatment labeled forks are stalled. More details can be found in the Material and methods section. **B-­-** HeLa S3 cells were labeled or not with EdU for 10 min (10’), or for 120 min in the presence of 1mM HU then subjected to an iPOND experiment. For the chase, cells were first labeled with EdU and 10 μM Thymidine was added for 120 min. EdU labeled DNA fragments were precipitated and co-­-precipitated proteins analyzed by western-­-blot for the presence of PCNA, KDM5A, KDM5B, RAD51 and H3 as a loading control. **C-­-** Proximity ligation assay (PLA) between KDM5A and PCNA in U2OS cells. Antibodies directed against KDM5A and PCNA were used either separately or together, as indicated. Representative images are shown. The number of spots per cell was counted in each condition using the Colombus software. Results are presented as box-­-plots showing the median, the 25 % and 75% quantiles and extrema below the images, Number of counted cells is > 100 cells for each point. * indicates a pvalue < 10^-­-35^ calculated by Wilcoxon test. **D-­-** As in **C**, except that KDM5A antibodies and ChK1 antibodies were used.

Given that KDM5A behaves like PCNA, we investigated whether it could be in contact with PCNA in cells by performing Proximity Ligation Assays (PLA). We observed a strong PLA signal only when the two relevant antibodies were included (Fig 4C). Importantly, this signal was preferentially detected in S-phase cells (Fig. S5A) and decreased upon knockdown of KDM5A (Fig. S6A), indicating that it is specific for KDM5A. Thus, these data indicate that endogenous KDM5A and PCNA interact. We next tested the proximity of KDM5A and Chk1, since KDM5A is important for ATR-mediated phosphorylation of Chk1 (see above). Moreover, Chk1 is a known partner of PCNA that localizes at fork in unchallenged conditions [43, 44]. We observed a PLA signal only when anti-KDM5A and Chk1 antibodies were mixed (Fig4D), and this signal was enriched in S-phase cells (Fig S5B), and decreased upon depletion of either KDM5A or Chk1 (Fig S6B). Taken together, these data indicate that in unchallenged conditions KDM5A is recruited to the replication fork in close association with PCNA and Chk1. Moreover, its recruitment is regulated by replication stress, like that of PCNA and Chk1, strongly suggesting that the effect of KDM5A on Chk1 phosphorylation by ATR is direct.

### KDM5A/B-mediated up-regulation of RRM2 is crucial for the acquisition of tolerance to replication stress

KDM5A is a major molecular driver of drug tolerance in cancer cells, allowing the generation of the so-called Drug tolerant persisters (DTPs; [17, 18]. On the other hand, resistance of cells to HU depends on the up-regulation of RRM2, likely to provide enough dNTPS to support repair of replication stress-induced DNA damages [45, 46]. Because our results indicate that KDM5A/B is important for the regulation of RRM2 and for RS-induced checkpoint, we postulated that KDM5A/B could be involved in the tolerance of cells to HU. To address this question, we generated U2OS cells resistant to HU by growing the cells in 0.25mM or 0.5mM HU until cells acquired resistance to the drug. Both concentrations led primarily to a block of cells in S-phase (data not shown). Cells quickly adapted to the presence of 0.25mM HU and a population of HU resistant cells was obtained after 10 days of treatment and called H25. In the case of 0.5 mM HU, a large proportion of cells died and a population of HU resistant cells was obtained following 1 month, and called H50. Both H25 and H50 showed a marked tolerance to increasing doses of HU, compared to U2OS (Fig 5A). This correlated with a lower activation of Chk1 (perhaps due, at least in part, to a decreased expression of Chk1, Fig. 5B) in response to 1mM HU in both cell lines compared to parental U2OS (Fig 5B). This lower activation is probably required for these cells to escape the S phase checkpoint. As expected, RRM2 was up-regulated several fold at both the mRNA and protein levels in both H25 and H50 cell lines compared to U2OS (Fig 5C and D). Although KDM5A mRNA levels were increased in H25 and H50 cells compared to U2OS, protein amounts were largely similar in all cell lines. By contrast, KDM5B was more expressed both at the mRNA and protein levels in H25 and H50 cells compared to U2OS (Fig. 5C and D). Thus, we conclude that the two cell lines we generated as tolerant to replication stress harboured up-regulation of KDM5B and RRM2.

**Figure 5:**
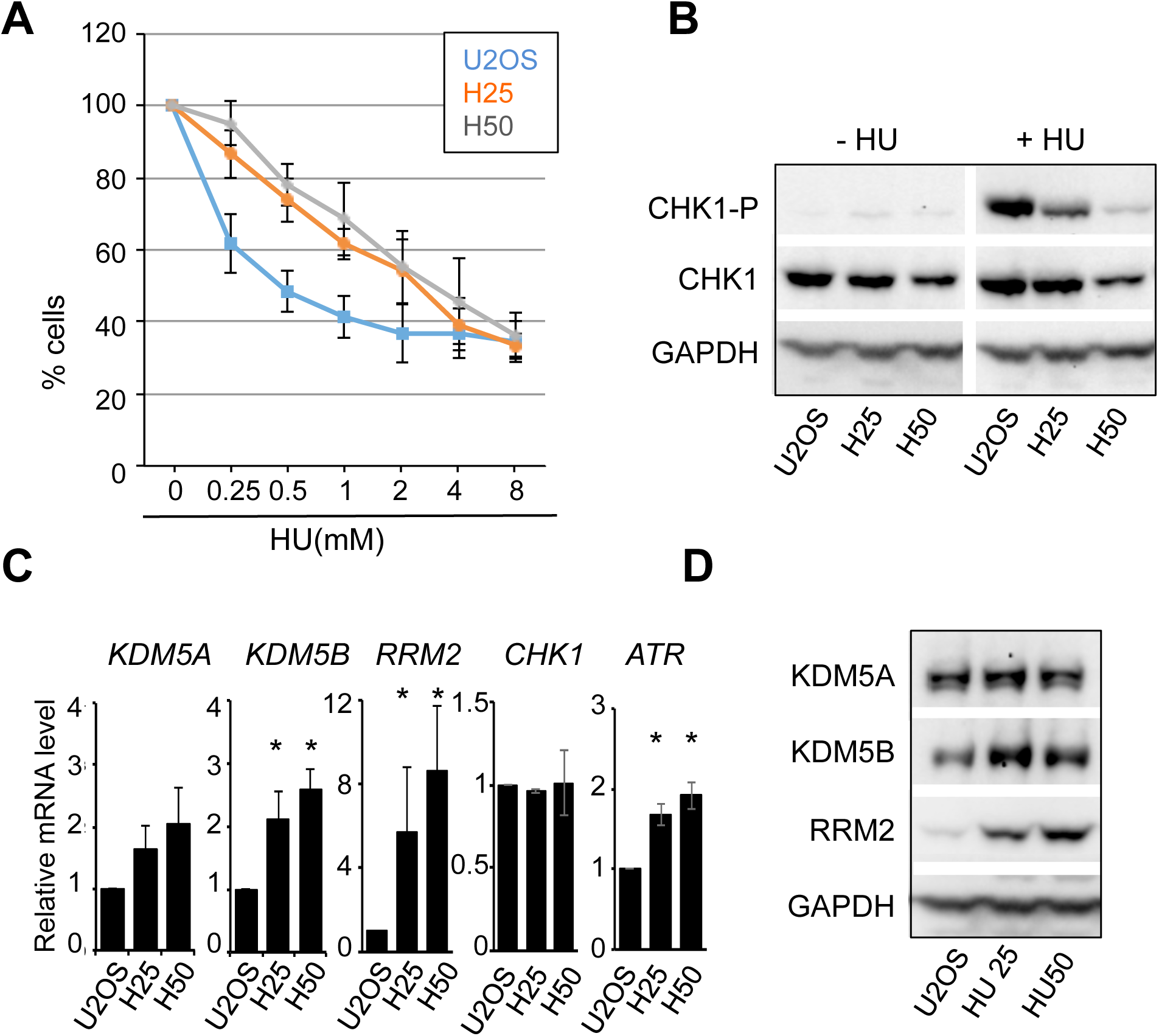
HU tolerant cells overexpress RRM2 and KDM5B. **A**-­-Viability of U2OS, H25 and H50, measured by WST assay, 72 hours following treatment with increasing doses of HU, as indicated. **B**-­-Western-­-blot analysis of CHK1 and S354-­-phospho CHK1 (P-­-CHK1), in U2OS, H25 and H50 cells before and following 1 hour treatment with 1mM HU. **C-­-** mRNA expression levels of KDM5A, KDM5B and RRM2, in U2OS, H25 and H50 cell lines. mRNA expression are normalized with the reference gene P0, and calculated relative to 1 for the siCtle. The mean and standard deviation from 3 independent experiments are shown. * pvalue<0.05 using a paired t-­-test. **D**-­-levels of KDM5A, KDM5B and RRM2 were analyzed by western-­-blot in the parental U2OS cells and its HU tolerant derivatives H25 and H50, grown in the presence of HU at 0, 0.25, and 0.5 mM respectively. GAPDH is used as a loading control. A representative experiment is shown.

These data are consistent with the possibility that this increase in KDM5B expression could be involved in RS tolerance through the up-regulation of RRM2. Because the two HU tolerant cell lines behaved similarly, we decided to work with the H50 cell line to address this question.

We first looked at the effect of depleting KDM5A and KDM5B on the tolerance of H50 cells to the continuous presence of 0.5 mM HU in the culture medium, and *per se* replication stress. Depletion of RRM2 was used as a positive control. Upon 48 hours depletion of KDM5A and KDM5B, the expression of RRM2 was diminished at both the mRNA and protein levels (Fig 6A and 6B), similarly to what we observed in U2OS cells. To evaluate the effect on the viability of H50 cells, we counted cells 72 hours following siRNA-mediated depletion of KDM5A/B or RRM2. In both cases, the capacity of cells to grow in the presence of 0.5 mM HU was reduced (Fig 6C). We next tested the involvement of KDM5A/B enzymatic activity in this function by treating cells with the KDM5 inhibitor CPI-455 or DMSO. As expected, treatment with CPI-455 led to an increase in H3K4me3 levels (Fig 6D). However, as in U2OS cells (Fig 1F-G), CPI-455 treatment did not decrease RRM2 expression at the mRNA or protein levels (Fig 6D-E). Accordingly, we found no effect on the survival of H50 cells following 72 hours of treatment with CPI-455, even if these cells were cultivated in the continuous presence of 0.5mM HU (Fig 6F). Thus, taken together, these data indicate that KDM5A/B expression, but not enzymatic activity, is required for the tolerance of H50 cells to replication stress.

**Figure 6.**
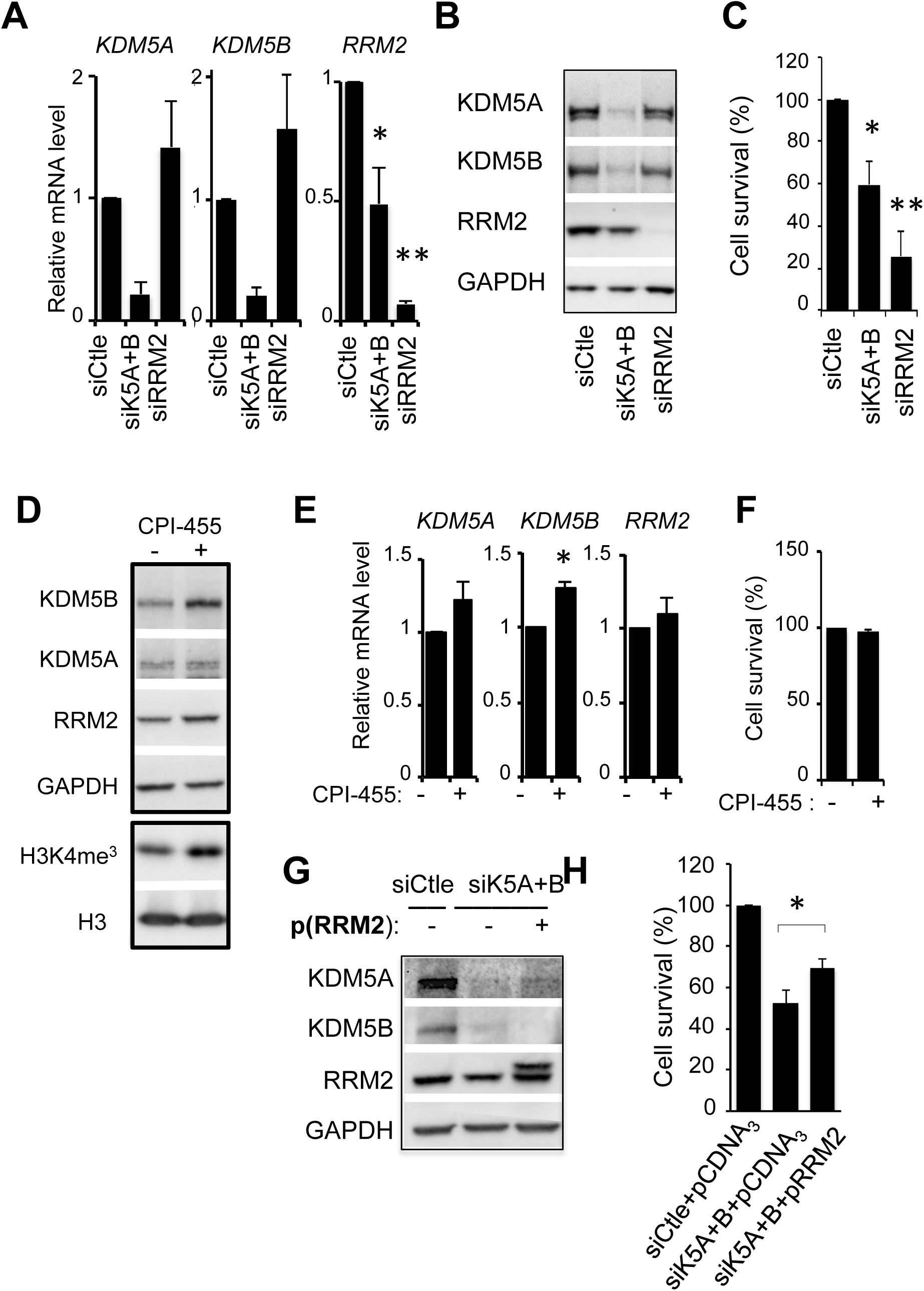
KDM5A/B are required for replication stress tolerance through regulation of RRM2. **A-** Relative mRNA expression levels of KDM5A, KDM5B, and RRM2, upon transfection of the indicated siRNA in H50 cells. Expression levels were normalized to the reference gene P0 and calculated relative to 1 for the siCtle sample. The mean and standard deviation from 4 independent experiments are shown, following normalization to 100 for siCtle treated cells. A paired t-test indicates significant difference between siK5A+B or siRRM2 and siCtle treated cells (*: pvalue<0.05, **: pvalue<0.01) **B-** Western-blot analysis of KDM5A, KDM5B, RRM2 and GAPDH as a loading control from H50 cells transfected with the indicated siRNA. **C**-Percentage of living cells following transfection of the indicated siRNA. The mean and standard deviation from 3 independent experiments are shown, following normalization to 100 for siCtle treated cells. A paired t-test indicates significant difference between siK5A+B or siRRM2 and siCtle treated cells (*: pvalue=0.013, **: pvalue=0.011)**D-** Western-blot analysis of KDM5A, KDM5B, RRM2, GAPDH, H3K4me3 and histone H3 from U2OS cells treated each 24 hours with 12.5 μM KDM5 inhibitor CPI-455 (+) or DMSO (-) for 48 hours. **E-** Relative mRNA expression levels of KDM5A, KDM5B and RRM2 in cells treated, each 24 hours, with 12.5 μM KDM5 inhibitor CPI-455 (+) or DMSO (-) for 48 hours. Expression levels were normalized to the reference gene P0 and calculated relative to 1 for the siCtle sample. The mean and standard deviation from 3 independent experiments are shown. The star * indicates significant difference between siK5A+B and siCtle treated cells (pvalue < 0.05, paired t test). **F-** Percentage of living cells following treatment of H50 cells each 24 hours with 12.5 μM CPI-455 for 72 hours (+) or DMSO (-). The mean and standard deviation from 3 independent experiments are shown, following normalization to 100 for DMSO-treated cells. **G-** H50 cells were treated with the indicated siRNA and 24 hours later transfected with pCDNA_3_-RRM2 p(RRM2) (+) or the empty vector (-). Expression of KDM5A, KDM5B, RRM2 and GAPDH were analyzed by western-blot 24 hours after plasmids transfection. **H**-Percentage of living cells 72 hours following electroporation of the indicated siRNA, combined to transfection of pCDNA_3_-RRM2 (pRRM2: +) or the empty vector (pCDNA_3:_ -) 24 hours later. To ensure efficient knockdown of KDM5A/B, cells were transfected once more with siRNA 24 hours following plasmids transfection. The mean and standard deviation from 4 independent experiments are shown, following normalization to 100 for siCtle/pCDNA_3_ transfected cells. * Statistical analysis with a paired t-test indicated that RRM2 surexpression significantly rescue the viability of K5A/B depleted cells with a pvalue <0.05.

We next tested whether this is due to their positive role on RRM2 expression. Strikingly, maintaining RRM2 expression by co-transfecting an expression vector for RRM2 in cells depleted for KDM5A/B (Figure 6G) partially restored the viability defect (Figure 6H), confirming that this defect is due, at least in part, to RRM2 down-regulation. All together, these results show that the tolerance of U2OS cells to HU depends on the up-regulation of RRM2, which is the consequence, at least in part, of KDM5A/B proteins overexpression.

## DISCUSSION

In this study, we show that KDM5A and KDM5B proteins are involved in the management of replication stress (RS), by regulating the level of RRM2, the regulatory subunit of the ribonucleotide reductase RNR, as well as by modulating the activation of Chk1 in response to RS. We further demonstrate the importance of the KDM5A/B-RRM2 axis in the resistance of cells to Hydroxyurea (HU), a potent replication stress inducer. Noticeably, the demethylase-activity of KDM5A/B is not required for this regulation and for tolerance of HU-induced replication stress.

### KDM5A/B is a positive regulator of RRM2

Our results show that KDM5A/B act as positive regulators of RRM2, the regulatory subunit of the ribonucleotide reductase (RNR). RNR is a tetrameric enzyme formed by the association of two RRM1 large subunits and two RRM2 small subunits. RRM2, but not RRM1, is regulated in a cell cycle dependent manner, by both transcriptional and post-transcriptional mechanisms. First, RRM2 is an E2F-target gene and thus its expression is increased in S-phase, a time when dNTPs levels need to be increased to allow DNA synthesis [30, 47]. Second, two pathways guide RRM2 to proteasome-mediated degradation: one operating in G2 phase depends on the Cyclin F-Skp2 ubiquitin ligase complex [48], the other operating in S-phase relies on APC/C^Cdh1^ [49]. Upon depletion of KDM5A/B, RRM2 expression decreases at both the mRNA and protein levels, indicating that KDM5A/B act primarily as a positive transcriptional regulator of RRM2, although a role on RRM2 stability cannot be excluded.

As an enzyme removing the H3K4me3 mark associated with active transcription, KDM5A is often described as a transcription repressor. Positive roles of KDM5A on gene expression have however been previously described although the precise mechanism is not clear. KDM5A associated to the MRG15 complex was proposed to favour the elongation step of transcription by demethylating H3K4me3 in the body of some genes, and as such in a reaction requiring its demethylase-activity [11]. Here, we show that KDM5A is mainly found at the promoter of RRM2, but not in the coding sequence and that the demethylase activity is not required, excluding such mechanism. A positive role of KDM5A expression was also observed on the expression of genes required for adipocyte differentiation. In a manner similar to what we describe here with KDM5B, KDM5A protein can be compensated by KDM5C in this process, and is found at promoter of genes that are regulated either positively or negatively by KDM5 proteins. However, by contrast to our study, in this case the demethylase activity of KDM5A is required for both types of regulation [28].

Interestingly, some examples in which KDM5A regulates gene expression or cellular processes independently on its enzymatic activity have already been described. The mechanism involved remains elusive but probably relies on interactions with regulators of gene expression. KDM5A was for example involved in the positive regulation of mitochondrial genes in drosophila, in a manner that depends on its PHD3 domain but not its demethylase activity. The authors proposed that the PHD3 domain allows the recruitment of KDM5A to chromatin by binding to H3K4me3, and may interact with transcriptional co-activators that remain to be identified [12]. In drosophila, Lid, which is the only expressed KDM5 family member, is crucial for larval growth in a manner that is independent of the JMJc domain carrying demethylase activity [50].

It is thus conceivable that KDM5A recruits transcriptional co-activators to the RRM2 promoter. Such a coactivator could be Tip60, a histone acetyl transferase contained in the MRG15 complex. Equally possible is the possibility that KDM5A impedes the access to the promoter or the activity towards chromatin of a co-repressor. Clearly, how KDM5A/B positively regulates RRM2, in a demethylase-independent manner merits further investigations.

### KDM5A localizes at forks and interacts with PCNA and Chk1

In this manuscript, we also provide data indicating that KDM5A plays a role at replication forks. Few studies described so far a role of KDM5 histone demethylases in the process of replication. KDM5C was shown to associate with PCNA and implicated in the assembly of the pre-replication complex by favouring the binding of CDC45 and formation of the active holo-helicase complex [40, 41]. Another study pointed to its role in the replication of heterochromatin together with Suv39h1 and HP1α, impeding transcription of heterochromatic region and genome instability [51]. Finally, KDM5A was shown to associate with ORC2 at centromeres to sustain genomic stability and prevent DNA-replication [52]. Here, we describe that KDM5A is enriched at active replication forks in close association with PCNA, and is required for full activation of Chk1. This ability to interact with PCNA is shared with KDM5C, shown to contain a PIP box of sequence QCDLCQDWF [40]. Interestingly, this sequence is conserved in KDM5A and likely mediates the binding of KDM5A to PCNA. We show by iPOND experiments that upon exposure to the replication stress inducer HU, KDM5A/B behave like PCNA and are released from the fork. Co-depletion of KDM5A and B leads to the inhibition of Chk1 activation, whereas ATR is properly activated and able to phosphorylate H2AX and RPA. This observation places KDM5A downstream of ATR and upstream of Chk1 in the replication stress pathway.

What could be the mechanism by which KDM5A controls Chk1 activation? It could be mediated through the transcriptional or post-translational regulation of key components of the activation of Chk1 by ATR (see Figure 3 and FigS3). However, given that KDM5A localizes at replication forks in a replication stress-dependent manner, it is tempting to speculate that it plays a direct role in Chk1 activation: For example, the capacity of KDM5A to be in close proximity to PCNA and Chk1 may favour the assembly of the complex required for activating Chk1 at forks, or may participate in Chk1 dynamics at replication forks. Indeed, Chk1 is bound to chromatin in unchallenged condition, and is released from the chromatin upon replication stress [53]. We reproduced this result but no change could be observed upon depletion of KDM5A/B (data not shown), suggesting that KDM5A does not regulate Chk1 binding to chromatin per se, but a step in its activation, which remains to be identified.

### KDM5A/B and replication stress tolerance

KDM5A is involved in the emergence of the so-called Drug tolerant persister cells (DTEPs) in cancer, in a manner that is dependent of its demethylase activity [17, 18]. This property, first described in PC9 cells treated with erlotinib was extended to other cellular models of cancers treated with distinct inhibitors either targeting components of signalling pathways or having a more general cytotoxic effect. In particular, KDM5A up-regulation allows the emergence of cells tolerant to cisplatin, a DNA damaging agent, in a manner dependent of its demethylase activity [18, 54]. Whether such a mechanism is conserved in cells becoming resistant to the replication stress-inducer hydoxyurea (HU) has never been reported. To answer this question, we derived from U2OS cell lines H25 and H50 that tolerate the presence of 0.25 mM and 0.5 mM HU, respectively. In contrast to what is described with cisplatin, we show that KDM5A levels do not change. However, KDM5B is up-regulated at both the mRNA and protein levels in H25 and H50 cells when compared to U2OS. Interestingly, KDM5B overexpression has been associated with chemotherapy resistance in epithelial ovarian cancer [23], with the development and progression of Glioma [22], and with the presence of a subpopulation of slow-cycling cells involved in long-term tumor maintenance in melanoma [55]. Thus, either KDM5A or KDM5B could fulfil the same function in cancer development/drug resistance, depending on the cell context. Accordingly, we show that KDM5A and KDM5B act redundantly in the regulation of RRM2, meaning that they can compensate each other. This observation is in line with other studies describing a redundant role of KDM5 family members on gene expression [8, 27, 28], or the up-regulation of either KDM5A or KDM5B depending on cancer cell types [20].

Although KDM5A/B are important to regulate Chk1 activation in U2OS cells in response to HU, this does not seems to contribute to the resistance phenotype of H50 cells. Indeed, in these cells, Chk1 activation is compromised despite an increase in the expression levels of KDM5B. This defective Chk1-dependent checkpoint signalling probably allows H25 and H50 cell to proliferate in the presence of a concentration of HU that normally stops U2OS cells in S-phase. We rather found that the KDM5A/B-dependent tolerance of H50 cells probably relies, at least in part, on the other mechanism we describe in our study, *i.e.* RRM2 expression control. This would fit with our finding that RRM2 is overexpressed in H50 cells and with the widely described function of RRM2 in allowing cells to cope with replicative stress.

Strikingly, although KDM5A/B expression is crucial for HU resistance, we report here that their histone demethylase activity is not required. Indeed, treatment with the KDM5 inhibitor CPI-455 had no effect on HU tolerance. On the contrary, it increases KDM5B and RRM2 in both U2OS and HU tolerant cell lines. This stands in contrast to what has been previously described for tolerance to other drugs [17, 18]. One explanation of this difference could be that previous reports addressed a role of KDM5A in the initiation of drug tolerance and not on its maintenance, which is clearly the step we study using the H50 model. Alternatively, the mechanisms involved may be drug-specific with specific mechanisms taking place to achieve tolerance to replication stress. As previously discussed, KDM5A and KDM5B may recruit to chromatin distinct chromatin regulators and/or inhibit the activity of negative regulators such as HDAC and as such may regulate nuclear events independently of their demethylase activity [15],[12]. Nevertheless, our study reveals the importance of assessing the requirement of the demethylase activity in KDM5A/B oncogenic functions, which may depend on cell types and cancer stages, as well as on the chemotherapeutic used to treat cancer. Our study also underlines the importance of designing new allosteric inhibitors of KDM5A/B, that impede partners association instead of inhibiting demethylase activity.

## MATERIALS AND METHODS

### Cell culture

U2OS and HeLa S3 cells were obtained from ATCC and cultured in Dulbecco’s modified Eagle’s medium (DMEM-5.5g/L glucose) plus 10% FBS. Medium was supplemented with 100U/ml penicillin, 100µg/ml of streptomycin (Gibco) and 1mM of Sodium Pyruvate (only for U2OS). H25 and H50 cells were established from U2OS cell line, by adding Hydroxyurea at a concentration of 0.25 or 0.5 mM in the medium. Cells were maintained in a 37°C incubator with 5% CO2.

### Antibodies

The following antibodies were used : anti-KDM5A (D28B10-Cell Signaling), anti-KDM5B (CL1147-Thermoscientific), anti-RRM2 (2G1D5-Cell Signaling), anti-CHK1 (2G1D5-Cell Signaling for western-blot; C9358-Sigma-Aldrich for PLA), anti-CHK1 phospho-Ser345 (133D3-Cell Signaling), anti-ATR (E1S3S-Cell Signaling), anti-ATR phospho-Thr1989 (GTX128145, GeneTex), anti-RPA (Subunit 9H8) (Santa Cruz SC56770), anti-RPA phospho-Ser4/Ser8 (A300-245A-M-Bethyl), anti-RPA phospho-Ser33 (A300-246A-T-Bethyl), anti-PCNA (CBL407-Millipore), anti-RAD51 (SC8349-Santa Cruz), anti-BRCA1(SC642-Santa Cruz), anti-H3K4Me3 (12209-Abcam), anti-H3 (1791-Abcam), anti-GAPDH (MAB374-Millipore), anti-γH2AX (Ser139)( 20E3-Cell Signaling or JBW301, Millipore), anti-53BP1 (NB100-304-Novus)

### Transfection/electroporation

2.10^6^ cells were electroporated with double-stranded siRNA to a final concentration of 2µM using an electroporation device (Lonza 4D Nucleofector) according to the manufacturer’s specifications. Alternatively, siRNAs were transfected with interferin at a final concentration of 20nM following the manufacturer’s instructions. The following siRNA were used: siGENOME non-targeting control smartpool #1 and #2 from Dharmacon (Horizon-Discovery) were used as control. siKDM5A-1 and siKDM5B-1 were siGENOME smart pool purchased from Dharmacon (Horizon Discovery). Other siRNAs were purchased from Eurogentec including siKDM5A-2: 5’-GGAUGAACAUUCUGCCGAAdTdT-3’, siKDM5AB-2, 5’-GGAGAUGCACUUCGAUAUAdTdT-3’, siRRM2 5’-UGAACUUC UUGGCUAAAUCUUdTdT-3’. Total siRNAs amounts were identical in all samples of each experiment using a mix 1:1 of control siRNA, a mix 1:1 of KDM5A or KDM5B or RRM2 siRNA and control siRNA, or a mix 1:1 of siKDM5A and siKDM5B.

Transfection of plasmids was done using the U20S-Avalanche reagent (Cambio), as described by the supplier. The plasmid pCDNA_3_-RRM2 was purchased from Addgene. Empty pCDNA_3_ was used as control. For rescue experiments, cells were first electroporated with siRNA and 24 hours later transfected with plasmids. siRNA transfection was done a second time at 48h using the Interferin^TM^ polyplus reagent (Ozyme). Cells were counted at 72 hours.

### Cell Viability and clonogenic assay

Cells viability was estimated either using WST assay or by counting the cells with trypan blue. For WST, cells were plated in 96 wells plate. After 24h, cells were incubated with or without HU or CPI455 (12.5µM). After 72h of treatment, WST-1[2-(4-Iodophenyl)-3-(4-nitrophenyl)-5-(2,4-disulfophenyl)-2H-tetrazolium] was added to the medium at a dilution of 1/10, followed by an incubation at 37°C for 2 hours, before measuring the absorbance at 450nm. Alternatively, viable cells were mixed with Trypan blue and counted using the countess II automated cell counter (Life technologies).

For clonogenic assay, U2OS cells were electroporated with siRNA as described before and seeded at 30 cells/cm2. The day after, they were treated for 24 hours with 50 μM HU or left untreated and then were allowed to grow for 10-15 days more before fixation and coloration with 1% Crystal Violet in H_2_O.

### Western-blotting

Cells were lysed with either Lysis buffer S (20mM Tris-HCl pH 7.2, 1% SDS) or N (20 mM Tris-HCl pH 8, 0.4% NP40, 300 mM NaCl, 1 mM DTT). Western Blots were performed using standard procedures. Antibodies used are listed in the paragraph “antibodies”. HRP-conjugated secondary antibodies were purchased from Amersham and Biorad.

### Total RNA extraction and RT-qPCR

RNA was extracted using an RNeasy mini kit (QIAGEN) as described by the supplier. 500 ng of purified RNA were reverse-transcribed by the PromII reverse transcriptase (Promega) using 0.5µg of random primers following the supplier’s protocol. cDNAs were analysed by q-PCR on a CFX96 real-time system device (Biorad) using the platinium SYBR Green qPCR SuperMix (Invitrogen), according to the manufacturer’s instructions. All experiments included a standard curve.

The primers used were **P0** forward : 5’-GCGACCTGGAAGTCCAACT-3’ and reverse 5’-CCATCAGCACCACAGCCTTC-3’; **KDM5A** forward **5’-**TGAACGATGGGAAGAA AAGG-3’ and reverse 5’-AGCGTAATTGCTGCCACTCT-3’; **KDM5B** forward 5’-GAGCTGTTGCCAGATGATGA-3’ and reverse 5’-TGATGCAGGCAAACAAGAAG-3’; **RRM2** forward 5’-TTCTTTGCAGCAAGCGATGG-3’ and reverse 5’-TTCTTTGCAG CAAGCGATGG-3’; **CLASPIN** forward **5’-**TAAACCACGGCTAGGTGCTG-3’ and reverse 5’-AGGCTTCCAGTTCTCTGTTGG-3’; **TOPBP1** forward 5’-AGCCCTCAACTG AAAGAGGC-3’ and reverse 5’-AACTCCACCTGTAATCTGCTCC-3’; **RAD9** forward **5’-** CTTCTCTCCTGCACTGGCTG-3’ and reverse 5’-CTTTGGCAGTGCTGTCTGC-3’; **RAD1 forward 5’-**CAGGGACTTTGCTGAGAAGG-3’ and reverse 5’-GGCCACAAGGCT GTACTGAT-3’; **MDC1** forward 5’-TCCGACGGACCAAACTTAAC-3’ and reverse 5’-ATCAGTGACCAGGTGGGAAG-3’; **HUS1** forward **5’-**CAGAAACGTGGAACACATGG-3’ and reverse 5’-ACAGCGCAGGGATGAAATAC-3’; **CHK1** forward 5’-AGAAA GCCGGAAGTCAACAC-3’ and reverse 5’-AGACTTGTGAGAAGTTGGGCT-3’; **ATR** forward **5’-**ACATTTGTGACTGGAGTAGAAGA-3’ and reverse 5’-TCCACAATTGGTG ACCTGGG-3’; **CDC6** forward 5’-GCAAGAAGGCACTTGCTACC-3’ and reverse 5’-GCAGGCAGTTTTTCCAGTTC-3’; **CCNE1** forward 5’-AGGGGACTTAAACGCCACTT-3’ and reverse 5’-CCTCCAAAGTTGCACCAGTT-3’.

### High-throughput microscopy

The Operetta automated high-content screening microscope (PerkinElmer) was used for quantification of gH2AX, 53BP1 bodies and/or cell cycle analyses. Cells seeded on 96 wells plate were fixed with 4% of freshly prepared paraformaldehyde and permeabilized with 1%Triton X-100 in PBS. A blocking step was performed with 1%BSA for 30min at room temperature. Cells were then incubated with primary antibodies in PBS-1%BSA overnight at 4°C. After three washes, cells were incubated with secondary antibodies (goat anti-rabbit Alexa 647 and/or donkey anti-mouse Alexa 488) at a 1/1000 dilution in PBS-1%BSA for 2h at room temperature. After three washes, a DAPI staining was performed for 10min. For labelling S-phase, cells were labelled with EdU for 20 min prior to fixation. EdU was revealed using the click-it imaging kit (Thermofisher) following the supplier’s instructions. Antibodies are described in the paragraph “antibodies”. Image acquisition with a 20× objective lens was automated to obtain at least 20 fields per well, allowing the visualization of a total of 500-1000 cells (three wells were acquired for each condition). Each picture was analyzed with the integrated Columbus software. Briefly, the DAPI-stained nuclei were selected (method B), and when necessary the size and roundness of nuclei were used as parameters to eliminate false positive compounds. For long term treatment with HU (50-100 mM for 24 h) the γ-H2AX staining was delineated using the find spot methods A or B and the sum intensity of the spots was measured. For short term treatment (1mM for 0-2h), the sum intensity of γ-H2AX per nucleus was measured. For cell cycle analysis, the sum of the DAPI intensity and the mean of the EdU intensity were plotted in order to separate G1, S, and G2 cells. The sum of the γH2AX intensity was subsequently determined in each of these cell population. When EdU labeling was not possible cells were separated in G1, S, and G2 phases according to DAPI sum staining. 53BP1 bodies were delineated using the find spot method B. Box-and-whisker plots of quantification of γH2AX staining were obtained with the R open source software R Core Team version 3.5.2 (2018-12-20; http://www.R-project.org/). They show the median, the 25 and 75% quantiles. Outliers, even if they are not shown are not excluded from the computations and tests (outliers are identified by not being in the range [25%Quantile−1.5 × InterQuantiles; 75%Quantile+1.5 × InterQuantiles] where interQuantiles=75%Quantile−25%Quantile). These representations have to be accompanied by statistical analysis of the comparison between the two populations. Statistical hypothesis tests were applied to confirm whether the hypothesis (that can be seen on the boxplot) that there is a differences between indicators of the two populations (such as mean, median, distribution) can be considered as true with a great confidence or can be due to random effect. Because data distribution was not normal (normality tested with Shapiro Wilk test), we used a Wilcoxon test to reject the hypothesis that the two populations medians are the same and thus conclude that there is a significant difference between the two medians if the *P* value is < 0.05, meaning a confidence of 95%.

### iPOND

We isolated proteins on nascent DNA (iPOND) as described previously [56, 57]. Newly synthesized DNA in Hela S3 cells (∼2.5.10^8^ per experiment) was labeled by incubation with 10 μM EdU for 5 minutes or for the indicated time in the presence of 1 mM HU. For pulse-chase experiments with thymidine (Sigma), cells were washed with cell culture medium supplemented with 10 μM thymidine, and incubated for 2 hours in thymidine-containing medium. Then the cells were crosslinked with 2% formaldehyde for 15 minutes. For the conjugation of EdU with biotin TEG azide, cells were permeabilized with 0.5% triton X-100, washed with 1xPBS, and then incubated for 2 hours in Click reaction buffer (10 mM Sodium-L-Ascorbate, 10 mM biotin TEG Azide (Glenresearch), 2 mM CuSO4). Cell pellets were washed with PBS, and then resuspended in lysis buffer (10 mM Hepes-NaOH, 100 mM NaCl, 2 mM EDTA, 1 mM EGTA, 1 mM PMSF, 0.2% SDS, 0.1% sarkozyl, Roche proteases inhibitor). Sonication was performed with a Misonix sonicator (fifteen cycles of 20 seconds sonication interspaced by a pause of 50 seconds). For the isolation of proteins on EdU - labeled DNA, samples were centrifuged 10 minutes at 18,000 x g and supernatants were incubated overnight with streptavidin-coupled magnetic beads from (Ademtech). An aliquot (2%) of the extract was kept as loading control. To reverse crosslinks and recover proteins bound to magnetic beads, the beads were washed in lysis buffer and then incubated in Laemmli buffer for 30 minutes at 95°C with shaking.

### Proximity Ligation Assay (PLA)

The in situ PLA was performed with DuoLink PLA technology probes and reagents (Sigma-Aldrich). Cells were fixed and processed as described above for immunofluorescence, except that the secondary antibodies were those provided with the PLA kit. Antibodies used are described in the paragraph “antibodies”. Revelation was performed according to the supplier’ instructions. Images were acquired with a fluorescence microscope (DM500, Leica) coupled to Metamorph and analysed using the Colombus program.

### Chromatin Immunoprecipitation

Cells were grown until 80% confluence and cross-linked with 2% formaldehyde for 10min before addition of 0.125M Glycine for 5min. Fixed cells were washed with PBS and harvested by scrapping. Pelleted cells were lysed with the following buffer: Pipes 5mM pH 8, KCl 85mM, NP-40 at 0.5%. The lysis was followed by homogenisation with a Dounce homogeniser. Nuclei were harvested by centrifugation and incubated in a nuclear lysis buffer: 50mM Tris pH 8.1, 10mM EDTA, 1% SDS. Samples were diluted ten times in a dilution buffer: 0.01% SDS, 1.1% Triton X-100, 1.2mM EDTA pH8, 16.mM Tris pH8.1, 167mM NaCl. A sonication step was performed ten times for 10s at a power setting of 5 and a duty cycle of 50% (Branson Sonifier 250) to obtain DNA fragments of about 500-1000bp. A preclearing step was made for 2hours at 4°C with 50µl of previously blocked protein-A and protein-G beads (Sigma) for 200µg of chromatin. Beads blocking was achieved by incubating the agarose beads with 200µg of herring sperm DNA and 500µg of BSA for 3h at 4°C. After preclearing, samples were incubated with antibodies specific for KDM5A (1ug/ml) or without antibody as negative control overnight at 4°C. Then, 50µl of blocked beads were added to the immune complexes for 2h at 4°C on a rotating wheel. Beads were washed once in dialysis buffer (2mM EDTA, 50mM Tris pH8, 0.2% Sarkosyl) and five times in wash buffer (100mM Tris pH 8.8, 500mM LiCl, 1% NP40, 1% NaDoc). Elution from beads was achieved by incubation in elution buffer (1%SDS, 100mM NaHCO3). Crosslinking was reversed by adding to samples RNase A (10mg/ml) for 30min at 37°C and incubating with 4µl SDS 10% overnight at 70°C. After 2h of proteinase K treatment, DNA was purified on a GFX column (GFX PCR kit, Amersham) and analysed by q-PCR.

## AKNOWLEDGMENTS

The authors wish to thanks members of DT’s team for helpful discussions. This work was supported by grants from “Toulouse Cancer Santé” to DT, the Ligue Nationale contre le Cancer to DT (equipe labellisée) and MV (Ligue Régionale de la Haute-Garonne).

## Authors contributions

SG and MV performed most experiments. VC performed many experiments of Figure 1. CR performed the iPOND experiment of Figure 4 and contributes to helpful discussions on the other results. SG, MV, VC, DT conceived and analysed experiments. MV wrote the manuscript with help from DT and SG. M-J P and J-S H, thanks to their expertise in Replication and CHK1 signaling, help in designing the experiments and analyse the results. KS initiated the project on KDM5A and E2F target genes that was the start of this work.

